# Analysis of Local Variability and Allostery in Macromolecular Assemblies using Cryo-EM and Focused Classification

**DOI:** 10.1101/365940

**Authors:** Cheng Zhang, William Cantara, Youngmin Jeon, Karin Musier-Forsyth, Nikolaus Grigorieff, Dmitry Lyumkis

## Abstract

Single-particle electron cryo-microscopy and computational image classification can be used to analyze structural variability in macromolecules and their assemblies. In some cases, a particle may contain different regions that each display a range of distinct conformations. We have developed strategies, implemented within the Frealign and *cis*TEM image processing packages, to focus classify on specific regions of a particle and detect potential covariance. The strategies are based on masking the region of interest using either a 2-D mask applied to reference projections and particle images, or a 3-D mask applied to the 3-D volume. We show that focused classification approaches can be used to study structural allostery, a concept that is likely to gain more importance as datasets grow in size, allowing the distinction of more structural states and smaller differences between states. Finally, we apply the approaches to an experimental dataset containing the HIV-1 Transactivation Response (TAR) element RNA fused into the large bacterial ribosomal subunit, to deconvolve structural mobility within localized regions of interest.

**Highlights:**

- Description of different image classification strategies in single-particle cryo-EM
- Quantitative evaluation of two classification methods using simulated data
- Application of the two classification methods to an experimental dataset

## 1. Introduction

Single-particle electron cryo-microscopy (cryo-EM) enables the visualization of macromolecules and their assemblies under near-native conditions [1]. In recent years, the technique has gained popularity, in part due to its ability to determine macromolecular structures at near-atomic resolution and without the need for crystallization [2]. While advances in resolution [3,4] have expanded the scope of the technique over the last five years, the ability to decipher structural heterogeneity is an ongoing area of development in the field [5,6]. Given that macromolecules, and especially their assemblies, are dynamic, image classification opens up the possibility to address novel types of questions pertaining to the molecular mechanisms underlying their function.

Structural heterogeneity can be either compositional or conformational in nature. Compositional heterogeneity means that the stoichiometry of subunits within an assembly varies within the dataset, such as particles containing or missing an additional, loosely associated protein factor. Conformational heterogeneity assumes that particles are uniform in composition, but the constituent components within each object can be flexible and can adopt one of several structurally different states. Conformational heterogeneity can be further subdivided into either discrete or continuous conformational heterogeneity. In the former case, the macromolecule would adopt one of several distinct structural states, each represented by a local minimum within the energy landscape describing all possible states. In the latter case, no distinct local energy minima exist, and the flexible regions can move in a mostly random manner. Finally, a fourth case can be defined as containing a combination of the above scenarios.

To understand structural heterogeneity within a single-particle experiment, the particle images are subject to a classification procedure, which assigns each particle to one of potentially many different classes. In the simplest scenario, a global classification strategy assigns each particle into a specific class on the basis of variability across the entire image. Different classification approaches have been developed, including supervised and unsupervised techniques, and numerous variations have been implemented to analyze structural heterogeneity [6-11]. Global 3-D classification does not require specific knowledge about the type and location of the heterogeneity, making it an integral part of today’s processing workflow of virtually all single-particle software packages. Given that macromolecular assemblies can be highly dynamic, and because every subdivision leads to fewer particles within each class (and thus lower signal and loss of resolution), the fundamental disadvantage of a global classification strategy is the limited number of well-defined classes that can be recovered from a dataset of a given size. This is particularly true when one wants to resolve variability in small, heterogeneous regions that may easily be lost during a global classification procedure. In contrast to a global classification strategy, “focused classification” zooms in on a region or feature of interest, in order to understand structural heterogeneity in a localized manner [12-15]. Focused classification can overcome the potential particle number limit associated with global classification by reducing the number of classes needed to represent the local variability and (in principle) excluding other regions of the particle from the analysis. This approach is particularly advantageous when regions outside of the area of interest are themselves dominated by structural heterogeneity. For example, minor domain movements within an otherwise dynamic macromolecular assembly might be difficult to resolve using global classification techniques alone because the majority of the signal guiding the classification procedure is dominated by regions outside of the area of interest. In another example, two large regions can exhibit independent variability, and a global classification may not converge on a solution that represents all possible states, or the number of states required leaves too few particles in the corresponding reconstructions, limiting their resolution. In general, focused classification provides an alternative means to deconstruct highly dynamic and/or heterogeneous datasets, reducing the analysis to a more tractable problem. Numerous successful applications of focused classification have been used to understand the independent movements of regions of large macromolecular complexes, such as the spliceosome and the ribosome [16-19].

Focused classification requires selecting a region of interest within the particle and excluding the remaining density. In the simplest implementation, a 3-D mask is applied to the reconstructed densities after each iteration to select the area of interest, and standard global classification is then performed using the masked reconstructions as references. A typical example of this is the classification of membrane proteins that contain detergent micelles: the 3-D mask is used to exclude the heterogeneous micelle while focusing on the protein [20]. The primary disadvantage of this “3-D masking” approach is that a projection of the density, which *only* contains the masked region, is compared with the particle image, which contains the masked region *in addition to* all other overlapping density, and this additional density can obscure the features to be classified. To reduce the problem of density discrepancy, the density outside the mask could be included in the reference after applying a low-pass filter [21,22]. The filter removes noise from the disordered regions of the particle while maintaining valid low-resolution signal to minimize the mismatch between reference and images. To further reduce density mismatch, another approach has been introduced, whereby, in addition to masking the 3-D object, the density outside the mask is computationally subtracted from the particle images [12,13,15]. This leaves a projection of the masked 3-D object and a density-subtracted 2-D particle image, which contains comparable features that can be used for classification. Another advantage of the “density subtraction” approach is that it can, in principle, be implemented in a hierarchical fashion, in order to subtract increasingly finer features in a step-wise manner. The (non-hierarchical) density subtraction approach has been used to improve heterogeneous regions of numerous macromolecular complexes that could not be improved using a global classification approach alone [12,13,15,19,23]. However, there are also disadvantages to this method. First, density subtraction requires an accurate measure of the signal in each particle image to properly subtract the desired density. Especially when looking at small regions and subtracting density corresponding to larger volumes, the subtraction may leave residual signal in the raw images, a problem that is exacerbated if the complex exhibits greater heterogeneity than is accounted for in the references used for density subtraction. The residual signal from the incomplete density subtraction can interfere with subsequent classification and obscure the variability in smaller regions (especially if applied in a hierarchical context). We and others have introduced another approach, where focused classification is performed in 2-D, with masks imposed on both the projection images and the experimental data [14,22]. In this alternative approach, a 3-D mask is defined for a region of interest, projected along the view determined for each particle and applied as a 2-D mask to the particle images and reference projections. Such an approach has been described in the context of bootstrap resampling and using the cross-correlation function to find the optimal solution [14] and has now been implemented within a likelihood-based framework in Frealign [8,22] and *cis*TEM [24]. The advantage of the “2-D masking” approach with focused classification is that it does not require signal subtraction, while constraining the classification to the area in the 2-D images that contain the region of interest and removing noise outside this region.

A major advantage of any focused classification approach is its ability to selectively classify features of interest within a distinct region of a cryo-EM map, which opens up numerous potential directions. First, it enables classification of pseudo-symmetric features in a particle that are related by a symmetry operator but not strictly symmetric due to independently dynamic mobility [15,25,26]. For example, surface-exposed regions of macromolecules may not obey the strict symmetry that may apply to the particle core, leading to loss of resolution in the surface regions of otherwise symmetric particles such as icosahedral viruses (reviewed in [27]). To classify pseudo-symmetric regions of a particle, the images are first aligned according to a common reference frame compatible with the pseudo-symmetry. The symmetry is then dropped, and multiple alignments for each particle image are determined, corresponding to all possible symmetry-related views, and an asymmetric reconstruction is calculated using each particle image multiple times to include all symmetry-related alignments. This effectively multiplies the number of particles in a dataset by the number of different possible symmetry operations and enables classification of different views into different classes, thereby resolving the heterogeneity in the pseudo-symmetric regions. This approach can, therefore, improve the resolution of density that would otherwise be an average of multiple structural states due to symmetrization. The approach has been applied, for example, to resolve density detail that was not visible after global classification alone [26], and to reveal genome structures within viral particles [28] (for other examples, see [27]). Second, selectively focusing on discrete asymmetric units can reveal covariant heterogeneity within the data. For example, two different regions located on opposite sides of a particle might be structurally coupled with each other. If the variability of two regions is random, there should be no correlation in the assignment of these regions to different classes during pseudo-symmetric classification. However, if correlation is present, this indicates covariance in the two regions. In the simplest case, counting of the number of matching asymmetric units within the same class, and comparison with a random distribution, would provide evidence for structural allostery. This phenomenon represents an area of development that may facilitate understanding global structural landscapes of dynamic macromolecular machines.

In this manuscript, we explore several different focused classification strategies with both synthetic and experimental data. We show the advantages and disadvantages of the “2-D masking” and “3-D masking” approaches, and additionally explore their ability to discover density covariances within otherwise distinct regions of a reconstruction. Finally, we show how focused classification can be applicable to heterogeneous experimental datasets, highlighting a particular test case that is relevant to visualizing mounted targets on scaffolds using single-particle cryo-EM.

## 2. Materials and methods

### 2.1 Generation of synthetic humanoid datasets

Synthetic datasets were generated as previously described [8]. Briefly, we randomly shifted and rotated projection images of humanoid structures, added noise, a CTF (to have CTF-modulated noise components), envelope function, and a final layer of noise. To reduce spurious correlations associated with the CTF for covariance analysis, we used a 640-pixel box size for projecting the data, and prior to the addition of noise and the CTF. 28 distinct datasets were made, corresponding to the different structural combinations of arms, hands, and feet (Figure 1). Combined datasets corresponding to the three distinct scenarios were then generated from the individual 28 datasets. Each combined dataset contained 10,000 particles (pixel size 5.24, box size 80 after Fourier resampling) with each of the 28 sub-datasets selected randomly.

**Figure 1 -.**
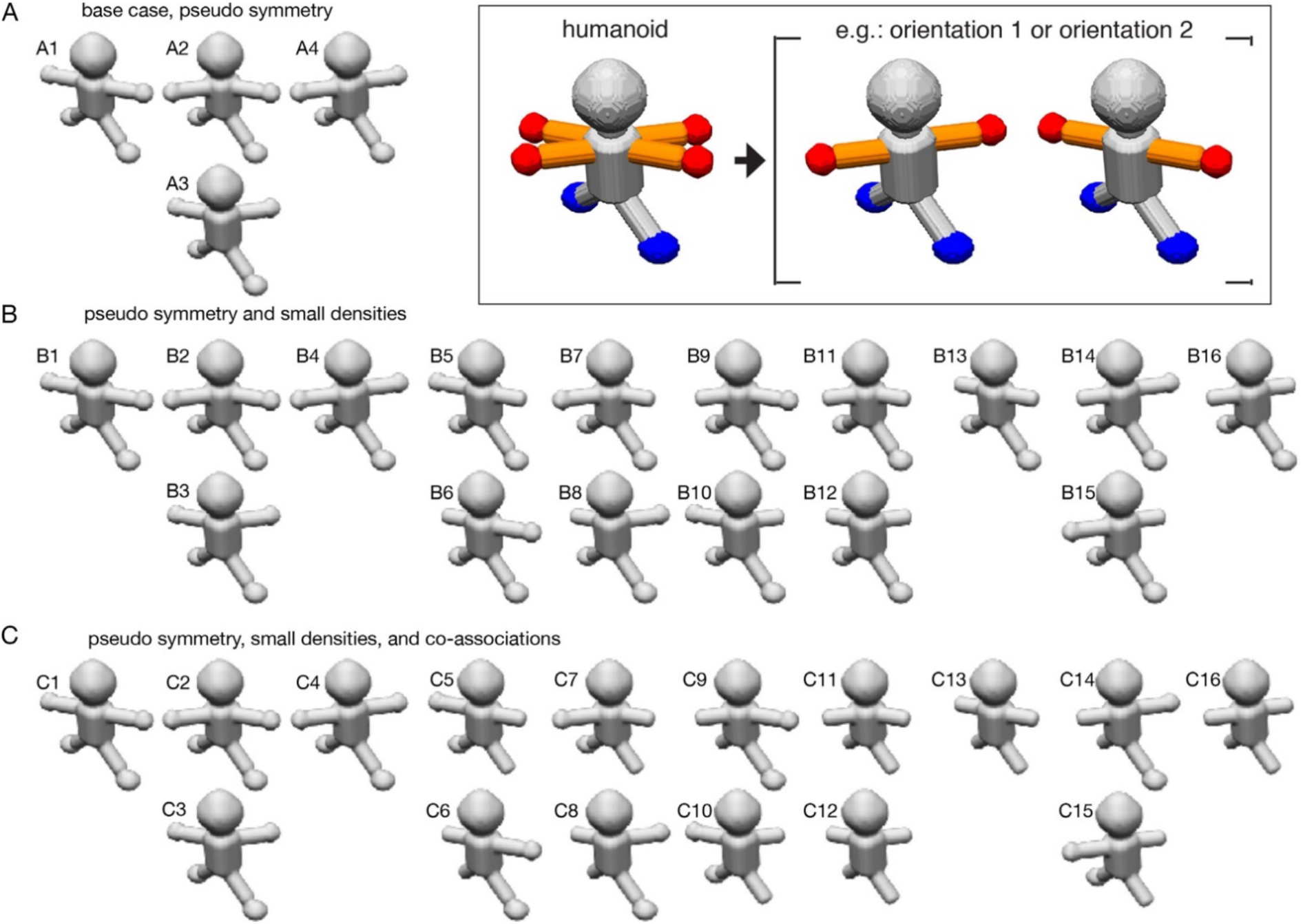
humanoid datasets and distinct scenarios used to assess focused classification. Different maps used to generate synthetic datasets described by the three scenarios are displayed. In each panel, A-C, two maps which are degenerate and related to one another by a 180° rotation are positioned vertically with respect to one another. The components used to generate the datasets are displayed in the inset, with the heterogeneous elements colored (arms, orange; hands, red; feet, blue). (A) For the base scenario, only the arms/hands are conformationally mobile. Four different combinations of maps lead to a dataset characterized by two different asymmetric units. Maps A2/A3 are related by a 180° rotation. (B) For the second scenario, in addition to the conformational mobility of the arms, the hands can be either present or absent. 16 different combinations of maps lead to a dataset characterized by four different asymmetric units. Maps B2/B3, B5/B6, B7/B8, B9/B10, B11/B12, and B14/B15 are related by a 180° rotation. (C) For the third scenario, in addition to the conformational mobility of the arms, the hands can be either present or absent, but their occupancy is *always* co-associated with a nearby foot. 16 different combinations of maps lead to a dataset characterized by four different asymmetric units. Maps C2/C3, C5/C6, C7/C8, C9/C10, C11/C12, and C14/C15 are related by a 180° rotation.

### 2.2 Particle assignment during focused classification

To facilitate quantitative assessment, we made the assumption that each classified particle belongs to the class with the highest probability (occupancy in Frcalign/*cis*TEM). At higher SNRs, this was an insignificant assumption, as most occupancies were close to 1; however, at lower SNRs, particles are represented by lower occupancies in multiple classes with slight differences between them. By assuming that each asymmetric unit corresponds to the class with the highest occupancy, we could simplify the calculation of κ coefficients and other analyses.

### 2.3 Measures for evaluating the accuracy of classification

To evaluate the accuracy of each classification trajectory, we define the following measures. For each asymmetric unit in each class:

- TP (true positive) — starting occupancy 100, ending marginal occupancy greater than all other classes.
- FP (false positive) — starting occupancy 0, ending marginal occupancy greater than all other classes.
- TN (true negative) — starting occupancy 0, ending occupancy less than the class with greatest marginal occupancy
- FN (false negative) — starting occupancy 100, ending occupancy less than the class with greatest marginal occupancy
- N: number of observations — TP+FP+TN+FN

Using the definitions above, the following metrics are defined:

Accuracy (the relative observed agreement among raters, or *Po*) = (TP + TN) / N
Sensitivity = True Positive Rate (TPR) = TP / (TP + FN)
Specificity = True Negative Rate (TNR) = TN / (TN + FP)
Kappa:

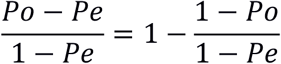

where *Po* is the accuracy, above, and *Pe* is the probability of chance agreement.
Youden’s Index (J Statistic) = TPR + TNR – 1.

### 2.4 Merging cryo-EM difference maps

Merging of the difference maps in Figure 4 was performed according to the following procedure. A merge volume was generated with 0s for the pixel values. Subsequently, for each pairwise difference map, and for each voxel, if the value of the voxel is greater than the value of this voxel in the merged map, set this as the value in the verged map.

### 2.5 Covariance analysis of separate regions of cryo-EM density maps

To determine whether different regions correlate with one another, normalized covariances were computed comparing fractional density occupancies of distinct components. An identical procedure was used for both scenarios 2 and 3. First, we performed 3-D focused classification, with the requested number of classes, k, identical to the expected number of non-degenerate asymmetric units. Binary masks were created for each region of interest (ROI), namely the hand in each of two positions, the near foot, and the far foot. The masks encompassed the ROI, with minimal incursion into neighboring density. A soft edge was not employed, because the mask was solely used for the purpose of computing fractional density occupancy values. For each of the k resulting maps, and for each ROI, the mask was used to extract the resulting density. Subsequently, the approximate mass in the ROI was calculated using the “volume” command implemented within the EMAN1 processing suite [29]. The resulting mass was optionally normalized to the true mass arising from a perfect classification to judge the quality of the classification, although this step is not strictly necessary for normalized covariance analysis. Finally, the normalized covariance matrix *R_ij_* was computed as:

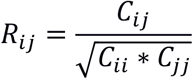

where *C_ij_* refers to the covariance between two components *i* and *j*. To make sure that there was adequate sampling, the resulting volumes represent an average of 3 independent runs, using random starting class occupancy values for initiating each classification.

### 2.6 Ribosome preparation

The 57-nt HIV-1 TAR element was appended inserted into twelve different helices (H9, H12, H19, H24, H25, H31, H45, H46, H59, H63, H68 and H98) by replacement of the loop residues to screen for optimal attachment sites. These twelve were chosen based on their location on the periphery of the ribosome and lack of tertiary contacts. All insertions resulted in viable bacterial growth (albeit much slower in some cases). H45 qualitatively yielded the most complete density with the least apparent mobility of the attached RNA (data not shown). Uniformly labeled ribosomes were prepared in the same way for all insertions. To ensure that all ribosomes contain the appended construct, a well-established protocol for introducing and characterizing site-specific mutations into *Escherichia coli* ribosomes was used [30,31]. Briefly, a *Δ7 prm E. coli* strain SQZ10 [32]. which has a genomic deletion of all rRNA genes, was used. The rRNA genes are supplied by a plasmid that also contains the *levansncrase* gene and confers kanamycin resistance (Plasmid 1, pHK-rmC-sacB). Levansucrase expression is lethal to *E. coli* when grown on sucrose-containing media [33]. An additional ampicillin-resistant plasmid containing the rRNA genes with the RNA construct of interest inserted (Plasmid 2, p278) was then transformed and grown in liquid culture. Cells were plated on media containing ampicillin and 5% sucrose to select for those that had lost Plasmid 1 but retain Plasmid 2. To confirm the selection, colonies were plated on Kan media to ensure that they cannot grow.

Insertion of TAR into helix 45 of p278 was carried out using site-directed ligase-independent mutagenesis [34]. Mutant plasmids were then transformed into SQZ10 cells and selected using the strategy described above. Mutant ribosomes were purified by first growing to mid-logarithmic phase (OD_550_ = 0.3-0.5) in 500 mL Luria Broth while shaking at 37 °C then chilled on ice for 30 minutes and pelleted by centrifugation. The cell pellet was then resuspended in 20 mL Resuspension Buffer (20 mM Tris-HCl, pH 7.5, 10 mM MgCl_2_, 100 mM NH_4_C1, 0.5 mM EDTA, 2 mM CaCl_2_, 6 mM β-mercaptoethanol). The resulting resuspension was lysed through a French Press three times, filtered through a 0.45 μm syringe filter and clarified by centrifugation at 18,000g for 30 minutes twice. The supernatant was concentrated to ~500 uL using a 50K MWCO filter (Amicon) and layered onto 36 mL 10-40% sucrose gradient in Gradient Buffer (50 mM Tris-HCl, pH 7.5, 10 mM MgCl_2_, 100 mM NH_4_C1, 6 mM β-mercaptoethanol) and ultracentrifuged in SW-32Ti rotor at 16,700g for 18.5 hours at 4 °C. 70S ribosomes fractions were collected, buffer exchanged into Storage Buffer (20 mM Tris-HCl, pH 7.5, 10 mM MgCl_2_, 100 mM NH_4_C1, 6 mM β-mercaptoethanol), aliquoted and stored at 4 °C until ready for grids.

### 2.7 Cryo-EM grid preparation and data acquisition

2.5 μl of purified ribosomes after sucrose fractionation were diluted to a concentration of 4 mg/ml with Storage Buffer and placed on UltrAuFoil Rl.2/1.3 300-mesh grids (Quantifoil) that were plasma-cleaned (75% argon/25% oxygen atmosphere, 15 W for 7 s using a Gatan Solarus). After 1 min incubation under >80% humidity at 4 °C, grids were blotted manually with a filter paper (Whatman No. 1) before being plunged into liquid ethane cooled by liquid nitrogen using a manual plunger. Leginon was used for automated EM image acquisition [35]. Grids were imaged on a Titan Krios microscope (FEI) operating at 300kV and equipped with a K2 Summit direct electron detector (Gatan). A nominal magnification of 22,500x was used for data collection, giving a pixel size of 1.31 Å at the specimen level, with the defocus range of −0.5 μm to −2.5 μm. Movies were recorded in counting mode with an accumulated total dose ~50 electrons/Å^2^ fractionated into 60 frames with an exposure rate of ~7 electrons/pixel/s.

### 2.8 Image processing and model generation

All pre-processing was performed within the Appion suite [36]. Motion correction was carried out by using the program MotionCor2 [37] and exposure-filtered in accordance with the relevant radiation damage curves [38]. The CTF for each micrograph was estimated using CTFFind4 [39] during data collection. 70S ribosomes served as a template for automatic particle picking using FindEM [40]. 346K particles were selected and subjected to per-particle CTF estimation using the program GCTF [41]. After 2D and 3D classification in GPU-enabled Relion [42,43], selected classes containing 232K particles were combined to a single stack and imported to Frealign for global refinement with 8 classes. Every ten cycles of refinement/classification, the reconstructed maps of all 8 classes were aligned to a common 50S scaffold using custom scripts implemented for performing a 3-D alignment within the Chimera package [44] while running Frealign/cisTEM, in order to maintain a common reference-frame for subsequent focused classification. A total of 50 cycles of global refinement/classification were performed. Subsequently, the best orientations were combined into a single parameter file for focused classification. Focused classification was performed for 500 cycles, and without further alterations to the orientations, by defining a spherical mask of 30 Å, centered on the expected region of TAR. Global resolution for the final map was estimated using the Fourier shell correlation (FSC [45]) at 0.143 and directional resolution anisotropy was evaluated by the 3D FSC server [46]. Local resolution estimation was performed using sxlocres.py implemented within Sparx [47].

The model of TAR attached to H45 of the 23S ribosome was prepared by first removing the loop residues of H45 from a recent 2.9 A structure, PDB ID 5AFI [48], and removing the polyA nucleotides from a model of TAR based on small-angle X-ray scattering data. The terminal backbone atoms were docked and aligned in UCSF Chimera [44]. The TAR region was then rigid-body refined into the cryo-EM density in Coot [49].

## 3. Results

### 3.1 Quantitative characterization of focused classification with 2-D and 3-D masking

3-D classification with different masking options, including the 3-D masking and 2-D masking, have been described and implemented within Frealign [8,22] and *cis*TEM [24]. In the present study, we quantitatively characterize the performance of these different options using simulated data, highlighting strengths and weaknesses of each approach. We generated multiple synthetic datasets that are characterized by various degrees of heterogeneity. Figure 1 shows the distinct components of a “humanoid” reconstruction, with the legs, body, neck, and head positioned identically, and representing the constant, homogeneous regions of a particle, characterized by twofold rotational symmetry. In contrast, the arms can belong to one of two conformations, and are therefore characterized by pseudo-symmetry. Lastly, the hands and feet, which represent small features of a map that might be lost during global classification, can be either present or absent. We generated maps representing all possible combinations of these features and created multiple synthetic datasets containing random translations and rotations, a contrast transfer function (CTF), an envelope function, and multiple levels of noise, bringing the final CTF-modulated SNR down to 0.100, 0,050, 0.025, 0.013, or 0.006, as previously described (Supplementary Figure 1 and [8,50]). Below, we describe three scenarios, which serve to demonstrate different aspects of focused classification. Importantly, in all described cases, focused classification is performed on an asymmetric subunit basis, which allows one to break down and constrain the heterogeneity problem [27] and reveal discrete movements within a more complex landscape of heterogeneity.

#### First scenario – the base, pseudo-symmetric case

In the base scenario, only the arms/hands are mobile and can adopt one of two distinct positions within an asymmetric unit, and the hand always remains co-occupied with an arm (Figure 1A). This case represents a common problem with pseudo-symmetric experimental datasets, whereby most of the molecule is homogeneous and characterized by symmetry (here, twofold), but one feature does not obey symmetry constraints (here, the arms/hands). There are four combinatorial possibilities, three of which would be expected to be recovered using a global classification strategy (structures A2 and A3 are degenerate and are related by 180° rotation). However, in an asymmetric focused classification centered on one side of the humanoid, one would expect to find only two non-degenerate possibilities, because the arm/hand can reside in only one of two structural states.

#### Second scenario – identifying small densities

In the second scenario, we use focused classification to recover finer features within a more complex structural landscape. In addition to the arms occupying one of two distinct positions, the hands can be either present or absent, and their occupancy is completely randomized (Figure 1B). Thus, for each of the four structural states described in the base scenario, one would see four additional structural states represented by the presence or absence of each of two hands. In sum, there are 16 different combinatorial possibilities, global classification would be expected to uncover 10 non-degenerate classes, but only four classes should be resolved using asymmetric classification.

#### Third scenario – identifying small densities and covariances

The third scenario is identical to the second scenario, except that a hand on each asymmetric unit is always co-associated with its corresponding foot (Figure 1C). For example, if the left hand is present, so is the left foot, and if it is absent, the foot too is absent; the same applies to the opposite asymmetric unit. One can then classify on the hand only, but look at both the hand and foot areas in the resulting maps and count the number of times that density for the hand co-occurs with density for the foot. In doing so, one can begin to decipher patterns and relationships within distinct components.

### 3.2 Focused classification on an asymmetric subunit of a synthetic humanoid

For each of the three cases described above, and for all five levels of noise, we performed focused classifications on a single asymmetric unit, with a mask around the region encompassing an arm and hand (Figure 2A). For these experiments, the particle alignment parameters were set to the correct parameters used to generate the data and were kept fixed during classification. To quantitatively evaluate the accuracy of classification, we used the κ coefficient as a statistical measure, which captures the performance of a diagnostic test, while taking into account the possibility of occurrence by chance [51]. We also used the Youden’s J statistic (informedness, [52]), but found that the results largely paralleled those of κ (data not shown). The κ coefficient evaluates the agreement of raters for classifying N items into mutually exclusive classes and relies on the precise knowledge of the number of false positives (FP), false negatives (FN), true positive (TP), and true negatives (TN), which we can obtain from the data (see Methods). Importantly, κ estimates the probability of an “informed” decision by taking into account random chance and returns 0 when classification is random (chance) and 1 when perfect classification is achieved. Qualitatively, it is simple to visually assess how “clean” the classification is, and whether or not the particles were correctly partitioned, by looking at the separation of the arms in our data. Supplementary Figure 2 shows how the results look when classification is nearly perfect (Supplementary Figure 2A), when classification is completely random (Supplementary Figure 2D), and two intermediate cases (Supplementary Figure 2B-C). A correct classification partitions the arms within a single asymmetric unit (and not its counterpart) into two distinct classes, with no signs of contaminating density (κ close to 1); as more errors are introduced, the two classes become progressively more mixed, up to a point where one cannot distinguish between the two volumes within or outside the asymmetric unit (κ close to 0, Supplementary Figure 2). In this manner, we could also determine which parameters provide optimal classification results (e.g. mask size, soft edge drop-off, etc., as demonstrated in Supplementary Figure 3), which we determined prior to evaluating the test cases.

**Figure 2 -.**
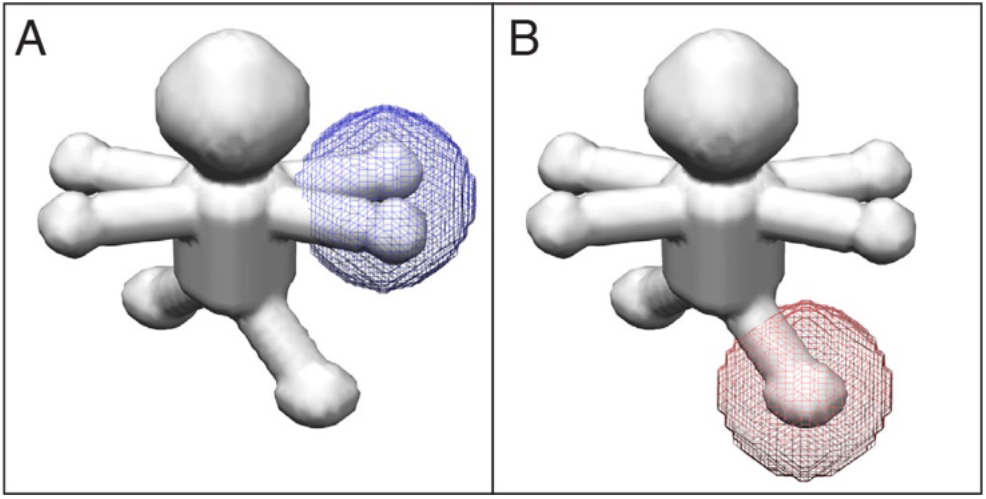
Application of masks onto regions of an asymmetric unit. Masks were applied either (A) onto the arm/hand region (blue) or (B) the leg region (red) prior to focused classification. Both types of asymmetric units are displayed, showing both orientations of the arm/hand combinations.

Table 1 shows the result of focused classification for all three scenarios, using both a 2-D masking approach and a 3-D masking approach, as implemented in Frealign and evaluated using the κ coefficient. The resulting numbers indicate the following general trends. First, for all three cases and for virtually all SNRs, the 2-D masking approach was superior to the 3-D masking approach. Such a result is not surprising because, as indicated in the introduction, the disadvantage of the 3-D masking approach, in the absence of density subtraction, is that the experimental projection images contain overlapping density along the path of the projection, as compared to a projection of the masked region from the reference map. The second general trend is that, with more mobile components within a dataset, and the smaller the desired features for detection, the lower the κ value and the more challenging it is to correctly classify the data. We observe major differences in accuracy between case 1 and either 2 or 3, because the latter contain more moving parts. However, the accuracies between cases 2 and 3 are roughly similar, likely because only small structural differences characterize the two datasets. Third, a lower SNR makes it more challenging to correctly classify the data, which is not surprising. However, it was surprising that, for the base scenario, even at the lowest SNRs and given how small of a feature we were trying to detect, we could still recover meaningful information and reasonably clean classes using the 2-D masking approach in particular, and to a lesser extent using the 3-D masking approach. In scenarios 2-3, higher SNRs were required to recover the correct classes (0.025 compared to 0.006, or ~4 times as high).

**Table 1.** – Results of focused classification on an asymmetric unit for the three different scenarios. Five different SNRs are evaluated, and the κ coefficient is displayed for the 2-D masking and 3-D masking case for each of three scenarios.

|  | Scenario 1 | Scenario 1 | Scenario 2 | Scenario 2 | Scenario 3 | Scenario 3 |
| --- | --- | --- | --- | --- | --- | --- |
| SNR | 2-D mask | 3-D mask | 2-D mask | 3-D mask | 2-D mask | 3-D mask |
| **0.100** | 0.99 | 0.91 | 0.85 | 0.70 | 0.87 | 0.71 |
| **0.050** | 0.96 | 0.85 | 0.73 | 0.56 | 0.75 | 0.61 |
| **0.025** | 0.87 | 0.74 | 0.42 | 0.41 | 0.46 | 0.39 |
| **0.013** | 0.72 | 0.57 | 0.21 | 0.17 | 0.21 | 0.17 |
| **0.006** | 0.47 | 0.36 | 0.09 | 0.08 | 0.08 | 0.09 |

Our experiments reveal that the 2-D masking approach, in its implementation within the likelihood-based framework of Frealign/*cis*TEM. does not completely isolate the area of interest from its surrounding density. While the 2-D masking approach produces more accurate results in the cases analyzed, its primary disadvantage is that projection images can contain additional density along the direction of the projection; if this density is homogeneous, it should be neutral in terms of classification, but if it is itself heterogeneous, it can bias the classification results. To account for this and to quantify the bias, we went back to the base scenario, where only the arm/hand combinations can move, but applied the mask onto an area of a leg and classified in that region (Figure 2B). We thus asked whether we can recover density for the arms, despite the mask being situated in a different location. As before, the number of correctly assigned particles was judged based on the arm/hand classes. If the arms completely determine the classification results, we would expect to see a κ coefficient of 1, whereas in the absence of crosstalk between arms and legs, the arms/hands would be randomly assigned and the κ coefficient would be 0. Table 2 shows that only at the highest SNRs does the heterogeneity outside of the area of interest influence the classification, and with a maximum κ coefficient of 0.23, the bias is not very severe. For SNR values of 0.025 and below, the results are effectively random. For the same dataset, a κ coefficient of 0.87 is obtained for an SNR of 0.025 when the mask is in its correct position around an arm. In contrast to the 2-D mask, when a 3-D mask is applied to the same location, the results are completely random at all SNRs. This is exactly what we would expect, because density outside this mask should not be introduced into a projection image after application of a 3-D mask. The above results indicate that bias generated by heterogeneity outside the area of interest is present but minor when using the 2-D masking approach, and absent in the 3-D masking approach.

**Table 2.** – Results of focused classification on an asymmetric unit when the mask is applied on the wrong region. Classification was performed after application of a 2-D mask or 3-D mask onto a leg (see Figure 2B), while the heterogeneity was characterized by the mobility in the arms/hands (scenario 1), and the κ coefficient was evaluated for the five SNRs and for each mask. Whereas the 2-D masking displayed some “leakiness” at the highest SNRs, the 3-D masking showed completely random classification.

| SNR | 2-D mask | 3-D mask |
| --- | --- | --- |
| **0.100** | 0.23 | -0.01 |
| **0.050** | 0.11 | 0.00 |
| **0.025** | 0.01 | 0.01 |
| **0.013** | 0.00 | 0.01 |
| **0.006** | 0.00 | 0.00 |
| **pure noise** | -0.01 | 0.00 |

### 3.3 Focused classification can identify covariant components in distinct regions of a map

Each individual object within a heterogeneous single-particle cryo-EM experiment can contain a unique combination of dynamic elements residing in distinct structural states. When multiple components are dynamic, and/or if they bind (or dissociate) in different regions, the conformational/compositional states of the components can be linked. Using focused classification, one can treat two distinct regions separately, and then ask whether there is any inter-dependence by calculating covariances within masked regions.

To evaluate covariance between distinct regions of a map, we used the datasets prepared for scenarios 2-3. In scenario 2, the presence of either hand, or either foot, are random and are not related to one another. In contrast, in scenario 3, the presence of a hand on one side of the humanoid is always correlated to the presence of a foot on that same side, whereas the opposite foot is randomly occupied and is not correlated to anything. Thus, one can apply a mask around the hands (encompassing both conformations), focus-classify the data, and then look for the presence or absence of a foot, which has not been subjected to focused classification. Quantitatively, once the dataset is classified and subdivided into groups, one would simply calculate the fractional density occupied by each component within the class (e.g. hand in position 1, hand in position 2, near foot, and far foot) normalized to its expected value, and compute a normalized covariance matrix (also known as a correlation coefficient matrix, see Methods) between the components. Since the presence of a foot is always correlated with the hand on the same side of the humanoid, irrespective of the conformation of the arm/hand, we further simplify the analysis by grouping both mutually exclusive hand positions into, more generally, a “near hand”. Thus, there are three regions for which fractional occupancies are computed – a “near hand” (blue in Figure 3), where the mask is applied for classification, a “near foot” (purple in Figure 3) on the same side of the humanoid, and a “far foot” (pink in Figure 3) on the opposite side of the humanoid. Given the nature of the mask, everything except for the hands is excluded from the classification. Since the mask is applied on an asymmetric-unit basis, the region that would otherwise constitute the “far hands” is not separated, and both mixed conformations are observed.

**Figure 3 -.**
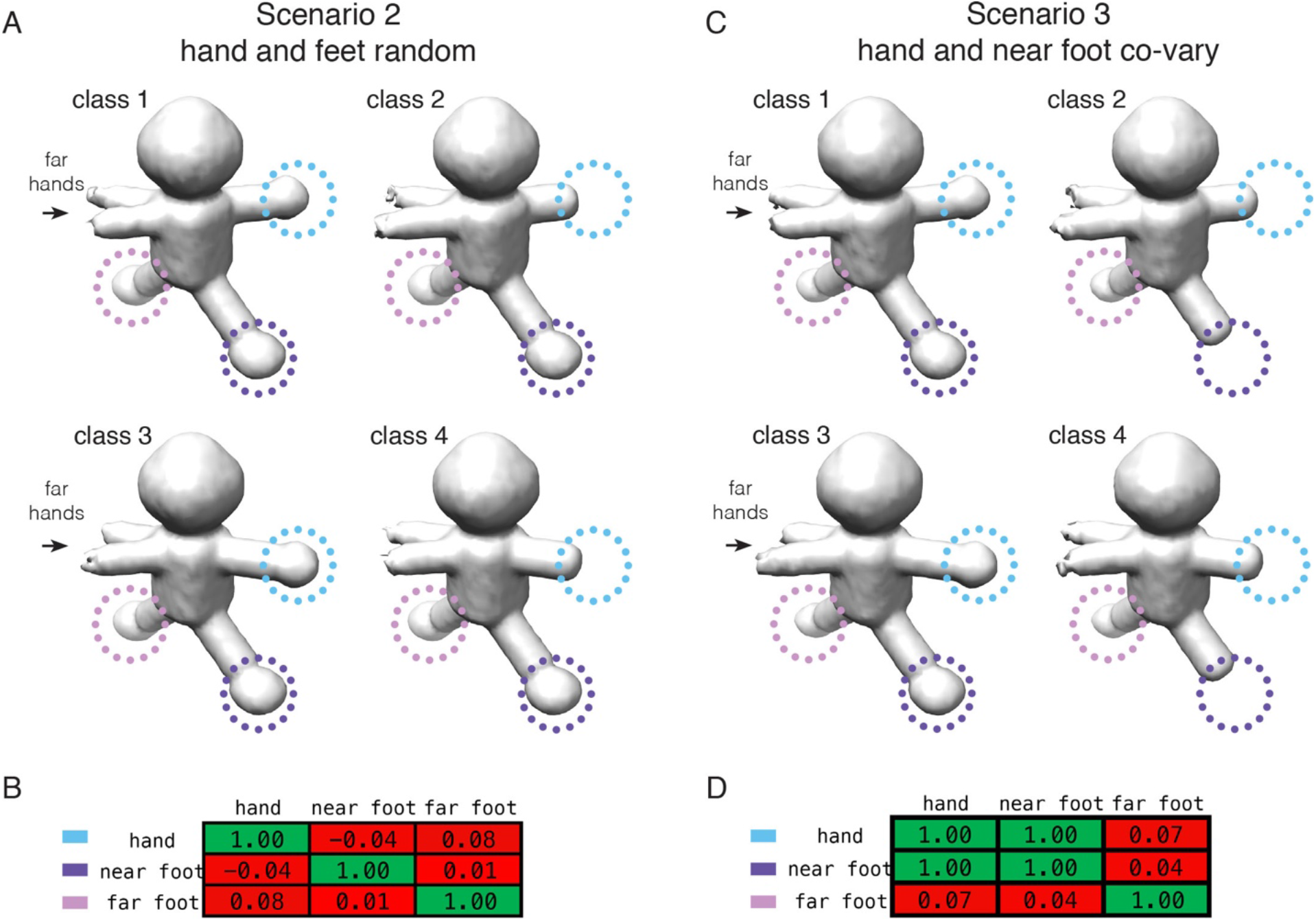
Evaluation of covariance within two different regions of a reconstructed object. Focused classifications using 2-D masks, applied to an arm/hand region (to the right of body in figure and encompassing both arm/hand conformations), were performed using either (A-B) the dataset for the 2^nd^ scenario or (C-D) the dataset for the 3^rd^ scenario, both at an SNR of 0.100. In all cases, four classes were recovered for the different asymmetric units (arms in two positions, each with and without a hand), and are displayed in panels A and C. (A) Volumes recovered from focused classification in the 2^nd^ scenario, where all components are randomly occupied (control). (B) Normalized covariance matrix describing the relationships between the near hand, near foot, and far foot. (C) Volumes recovered from focused classification in the 3^rd^ scenario, where the near hand is always co-associated with the near foot. (D) Normalized covariance matrix describing the relationships between the components. Near hand, where focused classification is performed, is circled in blue, near foot is circled in purple, and far foot is circled in pink. In the tables, the values are colored using a gradient: −1 (green, anti-correlated) < 0 (red, not correlated) < 1 (green, correlated).

For scenario 2, whereby no covariance is expected, the volumes captured through focused classification on an asymmetric-unit basis, and representing the four non-degenerate classes, are displayed in Figure 3A. As expected, they differ in the presence, absence, and overall conformation of the hands. For example, classes 1,2 or classes 3,4 differ by the presence or absence of a single hand; classes 1,3 or classes 2,4 either do or don’t have hands, respectively, but differ in the conformation of the arms; finally, classes 1,4 or classes 2,3 differ in both hand occupancy and arm conformation. Other than the hand/arm differences, no other regions of the maps have any apparent variability. Quantitatively, this is summarized by a normalized covariance matrix that describes the relative interdependence between the different components (Figure 3B). A value of 1 means that the pairwise occupancies of any two components are perfectly correlated, whereas a value of 0 means that they are completely random (a value of −1 means that they are anti-correlated). Identical components, related by the diagonal, are perfectly correlated, by definition. Otherwise, it is apparent that no two regions of the map are correlated to one another. This situation is different for scenario 3, however, which was designed to have the nearby hand and foot co-vary. The volumes captured through focused classification again represent the expected non-degenerate classes, and the hands/arms are related to one another in an identical manner as before. However, this time, it is clear that classes 2 and 4 are missing the nearby foot, whereas classes 1 and 3 maintain full occupancy. The normalized covariance matrix now shows that the hand is always co-associated with the nearby foot. The occupancy of the far foot, on the other hand, remains random, and is accordingly associated with a low normalized covariance value. The same experiment can be performed for more complicated combinations of hands and feet, but the principle is the same – that assessing the inter-dependence of density occupancies within distinct regions of a macromolecular complex can provide insight into hidden allostery within the data.

### 3.4 Focused classification facilitates deconvolving heterogeneous regions within an experimental dataset

The techniques described here have been used to decipher both conformational and compositional heterogeneity within biological samples (for example, [16,26,53]). In addition to the published results, one area where they will be particularly useful is to deconvolve conformational heterogeneity when using scaffolds for the purpose of structure determination. Several groups have shown that larger protein and/or nucleic-acid scaffolds can be used to aid in the determination of smaller structures, which by themselves would be too challenging to analyze [54,55]. However, the problem with all current approaches is that the particles of interest are not necessarily rigidly bound. Thus, the regions closer to the site of attachment will be characterized by less heterogeneity (and a lower B-factor), whereas the regions further from the site of attachment will exhibit more heterogeneity (and a higher B-factor). To demonstrate this, we used a bacterial 70S ribosome as a scaffold, and engineered in a fusion RNA representing the HIV-1 Transactivation Response (TAR) element. Subsequently, we performed either global classifications on the entire dataset or focused classifications on the region around TAR.

The HIV-1 TAR element was uniformly inserted into Helix 45 of the E. coli large 23S ribosomal RNA. Ribosomes containing the TAR knock-in were selectively purified (see Methods) and subjected to single-particle cryo-EM analysis. We collected 929 micrographs, providing 346,851 particles in the dataset (Supplementary Figure 4A). A single-model refinement, in the absence of any classification, showed high-resolution in the ribosome core, and lower resolution in the regions characterized by structural heterogeneity (Supplementary Figure 4B-C). Due to a large amount of mobility, the site of TAR fusion was only partially visible at the normal thresholds used for displaying the coulombic potential map. We then performed a global classification of the data, using a soft-edge spherical mask. This procedure resulted in distinct classes, separated according to the expected heterogeneity associated with purified bacterial ribosomes [56] (Supplementary Figure 4D). The combined differences are summarized with a merged map, demonstrating the full extent of heterogeneity for the global classification case (Figure 4A); notably, the resolved heterogeneity did not improve the density at the site of fusion. Subsequently, we performed a focused classification of the data using 2-D masks, applying the mask to the area where TAR has been inserted. As expected, the resulting maps were able to clearly separate out some of the different conformations of TAR (Supplementary Figure 4E). However, the majority of the normal ribosomal heterogeneity was largely ignored, as summarized by the merged difference maps (Figure 4B) and an overlay of the reconstructed classes (Figure 4C). In terms of characterizing classification performance, this result is important for several reasons. First, even though the area of interest is small, the focused classification approach using 2-D masks can partially deconvolve the density. Second, despite the extensive “normal” structural heterogeneity present on bacterial ribosomes (e.g. Figure 4A), which may confound the 2-D focused classification approach (e.g. Figure 2 and Table 2), we do not observe this in our results. We also performed focused classifications using 3-D masks, but the quality of the reconstructed TAR region was noticeably poorer (data not shown), consistent with the poorer performance of the 3-D masking approach using synthetic data (e.g. Table 1). These experimental results further demonstrate the ability of the 2-D masking approach to separate out local structural variabilities in the context of otherwise extensive global structural differences.

**Figure 4 -.**
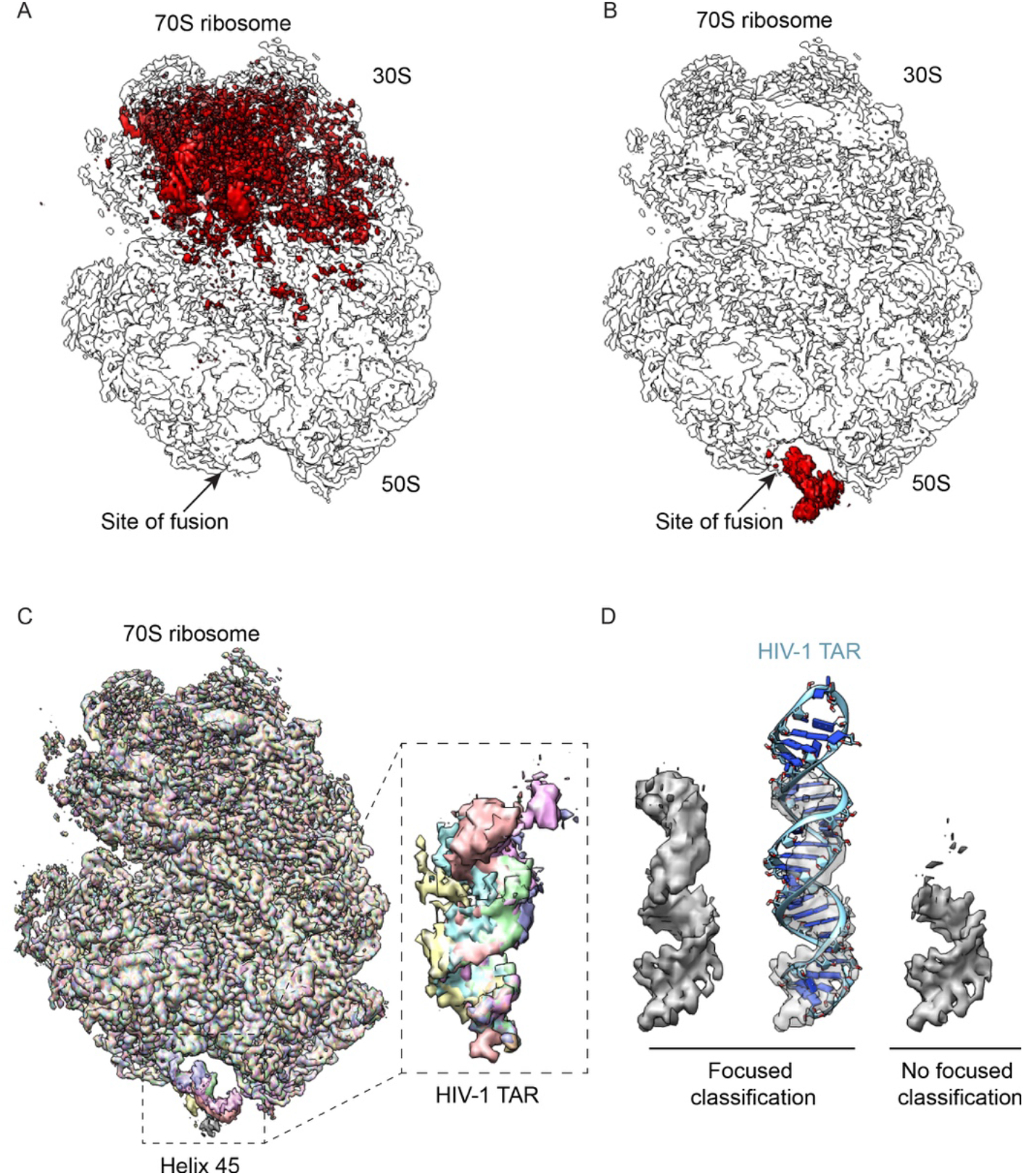
Experimental reconstructions highlighting the use of focused classification to analyze highly heterogeneous datasets. Bacterial 70S ribosomes containing an HIV-1 TAR element fused into Helix 45 (H45) were used to analyze different classification approaches. (A) Combinatorial pairwise differences between all 8 classes from global classification merged into a single volume to highlight the overall variability. (B) Same as A, but from the result of focused classification using 2-D masks, applied on the region of TAR fusion into H45. In both A-B, arrows denote the site of fusion. (C) Overlaid reconstructions after focused classification, highlighting the differences within the TAR element, but not in the rest of the ribosome. (D) Close-up of TAR reconstruction after deconvolving its mobility through focused classifications (left), shown also with a rigid-body docking of the TAR element into density (middle). A control reconstruction, without focused classification but using the same number of particles, is displayed alongside (right).

The best reconstruction of HIV-1 TAR showed a clearly defined RNA helix, a marked improvement over a global classification strategy alone (Figure 4D). The density was characterized by progressively poorer resolution, as a function of distance from the site of attachment. For a largely A-form HIV-1 TAR RNA helix, the behavior of the fusion can be thought of as a lever pivoting around a fulcrum; the further out from the point of attachment, the more inherent mobility, and thus the lower the resolution. A similar behavior has been observed with other scaffolding strategies, whereby the peripheral regions are characterized by lower resolution [54,55]. In addition to providing novel biological insight, focused classifications can broadly facilitate scaffolding approaches for solving structures of small proteins and RNAs.

## 4. Discussion

Using a synthetic dataset, we describe a quantitative assessment for several focused classification implementations within the Frealign/*cis*TEM processing packages. The algorithms have been used to classify features in several experimental studies [16,26,53], and we further demonstrate the applicability of the approaches for deconvolving heterogeneous regions within small scaffolded RNAs to facilitate the development of substrate supports for cryo-EM [54,55].

The present study will help users decide which strategy to use in a particular case. Focused classification using 2-D masks can be applied to individual asymmetric features (also known as symmetry expansion [27]), and, as implemented within Frealign/*cis*TEM. have generally been found to perform better than 3-D masking approaches, due to density mismatch between particles images and reference projections after 3-D masking. A possible disadvantage of the 2-D masking approach arises from the projection nature of the data. Any area within a 2-D projection image will not only contain density relevant to the region of interest, but also residual density along the projection path. If the residual density is itself heterogeneous, it can potentially confuse or bias the classification procedure (especially if the variability within the region of interest is significantly smaller compared to variability elsewhere). In Table 2, we demonstrate that this effect is real, at least with high SNR data. However, in practice this problem appears to be small, based on the results obtained with the synthetic data (compare Tables 1 and 2), and in an experimental setting in the context of large-scale global heterogeneity in the current work (Figure 4A-B), and in previous biological studies [16,18]. Conflating heterogeneity along the projection path would be treated as noise, in a manner that is perhaps analogous to incomplete density subtraction.

Our tests with the synthetic dataset demonstrate that additional questions, such as those pertaining to structural allostery, can be addressed in single-particle experiments. We showed how classifying variability in a region of a density map can reveal covariance with a secondary region, in this case between a hand and a foot. With synthetic data, such analyses are predicated upon having knowledge of the real density; in an experimental setting, an analogous approach would mask out regions corresponding to, for example, known components prior to analyzing the resulting normalized covariance matrices, as has been previously shown in one simplified example with ribosome-associated factors [57]. In general, the ability to classify independently on separate regions of a map provides opportunities to inter-relate distinct regions of an object beyond simply recovering densities, a form of computational identification of allostery within a system. Some cautions should be taken in the analyses of covariance. First, to avoid under-sampling, it is advisable to compute an equal or greater number of classes than expected. Second, and related to the previous point, classifications should be run multiple times, starting from different random particle seeds. Both of these precautions will ensure that sufficient pairwise occupancies have been calculated to reach statistical significance and avoid spurious correlations. Third, some caution should be taken in the interpretations of results using 2-D masks (due to the possibility of “leaky” biases during classification), although our experimental observations suggest that the biases should be minimal (Figure 4B). Finally, global classifications can also be used for the purpose of covariance analysis, and they can have specific advantages, as they would recover non-degenerate differences that are lost during classification on an individual asymmetric unit (which is easily seen with the experimental setup of the humanoid, as the number of non-degenerate structures (globally) far outnumbers the number of distinct asymmetric units). Whereas focused classifications help constrain the number of different classes and can simplify the analysis, the results should ideally relate to the global context of heterogeneity. In the future, more elaborate methods could be devised for broader applicability beyond pairwise covariances.

Our results using HIV-1 TAR fused to bacterial large ribosomal subunits show how focused classifications can help computationally deconvolve highly mobile features within experimental cryo-EM datasets. These data are particularly applicable for the development of structural scaffolds for the analysis of small proteins and RNAs [54,55]. The TAR fusions are universally mobile about a central fulcrum point, which corresponds approximately to the site of attachment, and the density is lost in the absence of proper classification. However, careful application of masks during focused classification enables partial recovery of some of the structural elements within the TAR fusion, visualizing most of the A-form RNA helix. Scaffolding approaches are gaining popularity in single-particle analysis, because small proteins may not have sufficient signal for accurate assignment of Eulerian orientations. Focused classification can help ameliorate problems associated with structural mobility and bring out the most of the structure of interest.

## Acknowledgements

We thank Dr. Kurt Fredrick for assistance and helpful discussions regarding the ribosome purifications, Bill Anderson for help with cryo-EM data collections. We acknowledge the support of NIH grants RO1 GM065056 (to KMF) and P50 GM103368 (HIVE Center, to KMF and DL). DL also acknowledges the support of DP5 OD021396. WAC was supported by a Pelotonia Postdoctoral Fellowship from the Ohio State University. NG is an Investigator of the Howard Hughes Medical Institute.

## Author Contributions

WAC, KMF, NG and DL designed the study. NG is the primary developmer of Frealign and cisTEM. DL prepared and performed calculations with the simulated data. CZ, WAC and YJ performed the calculations with experimental data. CZ, WAC, NG and DL analyzed the data. DL wrote the paper, with help from all authors.

## Code availability

Frealign and *cis*TEM are open-source and distributed under the Janelia Research Campus Software License (http://license.janelia.org/license/janelia_license_1_2.html). All scripts and datasets for these studies will be made available from the Lyumkis laboratory upon request.

---

**Supplementary Figure 1 -.**
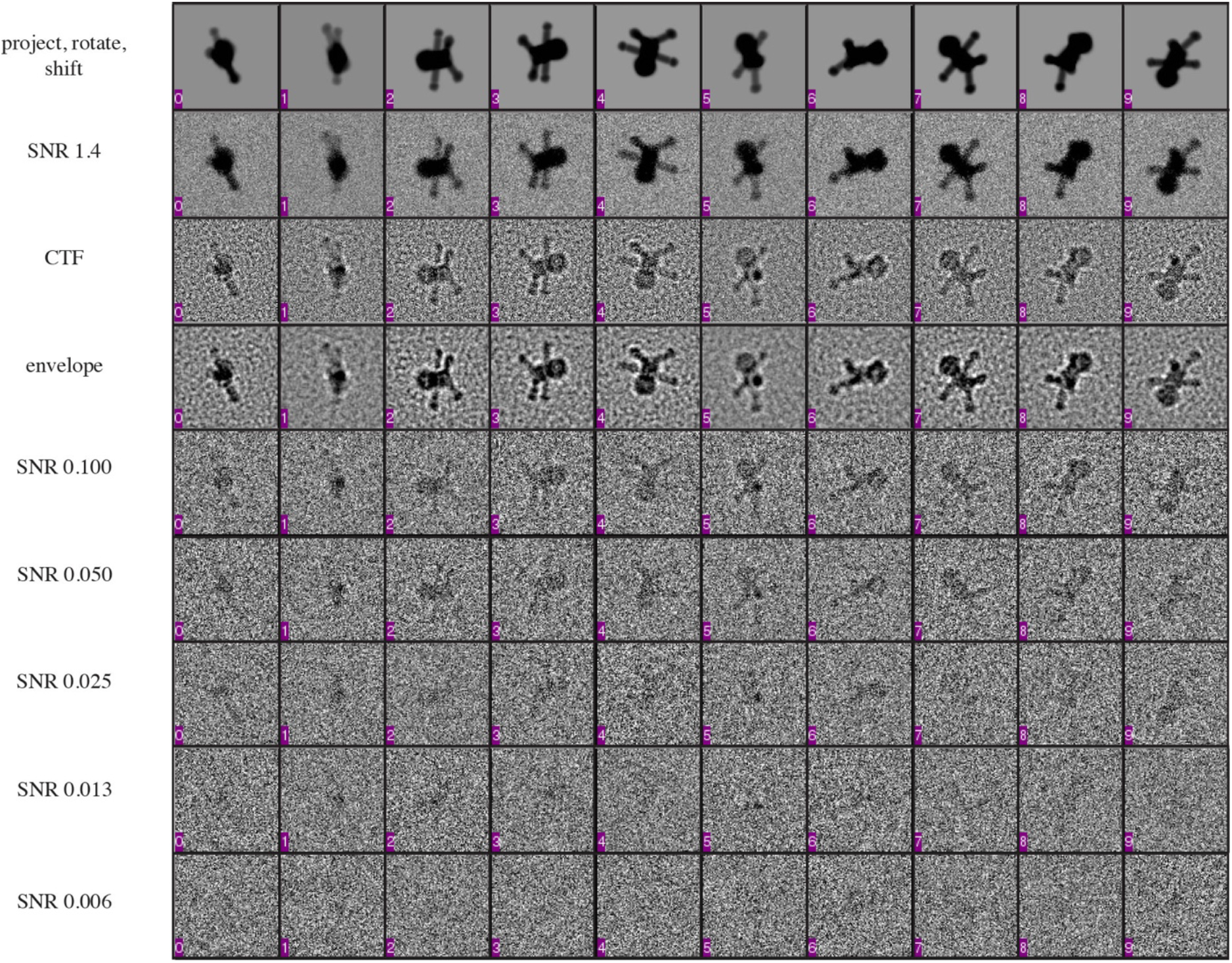
synthetic data generated from the humanoid volumes. Each volume was randomly projected, rotated, and shifted. Noise was then applied to the projection images, followed by a CTF and envelope function, and lastly the level of noise was brought down to one of five different levels (0.100, 0.050, 0.025, 0.013, and 0.006), as previously described [8]. The different projections were then randomly inserted into a 10,000-particle dataset for focused classification experiments.

**Supplementary Figure 2 -.**
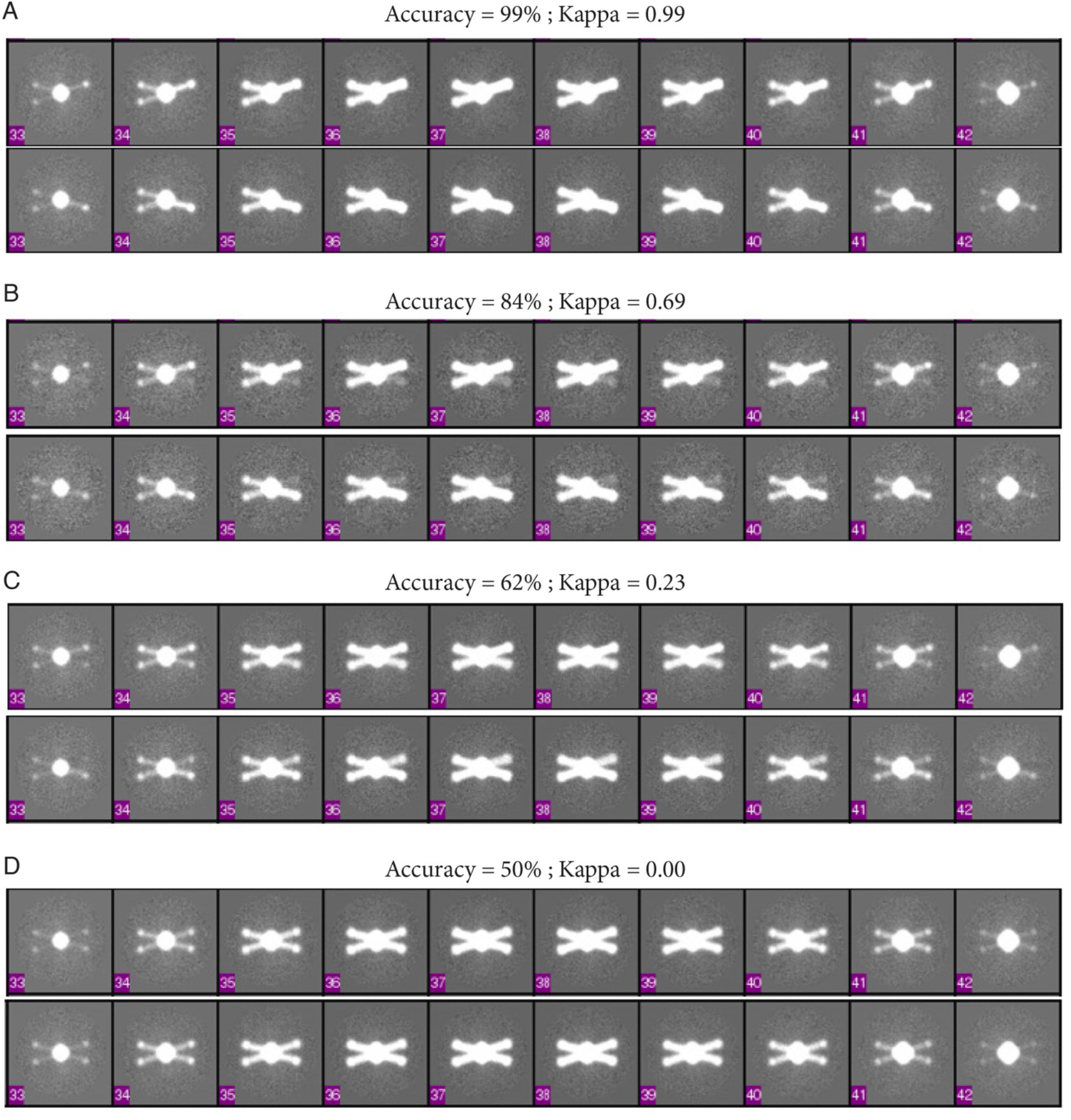
visual demonstration of classification accuracy. Slices through a reconstruction are displayed for each panel (middle slices 33-42 within a 96-slice volume, for each of two distinct classes [top and bottom]) around the Z-height of the arms. Classification was performed on the right asymmetric unit and for the base dataset, where two different classes are expected. Ideally, only the right arms would partition into one of several different classes. Classification was performed under four different levels of noise, which resulted in distinct accuracies. Panels A-D demonstrate how the accuracies, the associated κ coefficient, and the density varies with increasing errors. (A) Accuracy is nearly perfect, κ is close to 1 and the two classes show complete distinction in the right arm region. (B) Accuracy is worse, κ is has dropped to 0.69, and some contamination is evident in the opposing arm. (C) Accuracy has dropped further, κ is close to 0, and the two volumes become virtually indistinguishable, although some differences within the density amplitude point to residual heterogeneity. (D) Accuracy is completely random (50% represents a coin toss when two possibilities are present), κ is correspondingly 0, and no difference in the maps is evident.

**Supplementary Figure 3 -.**
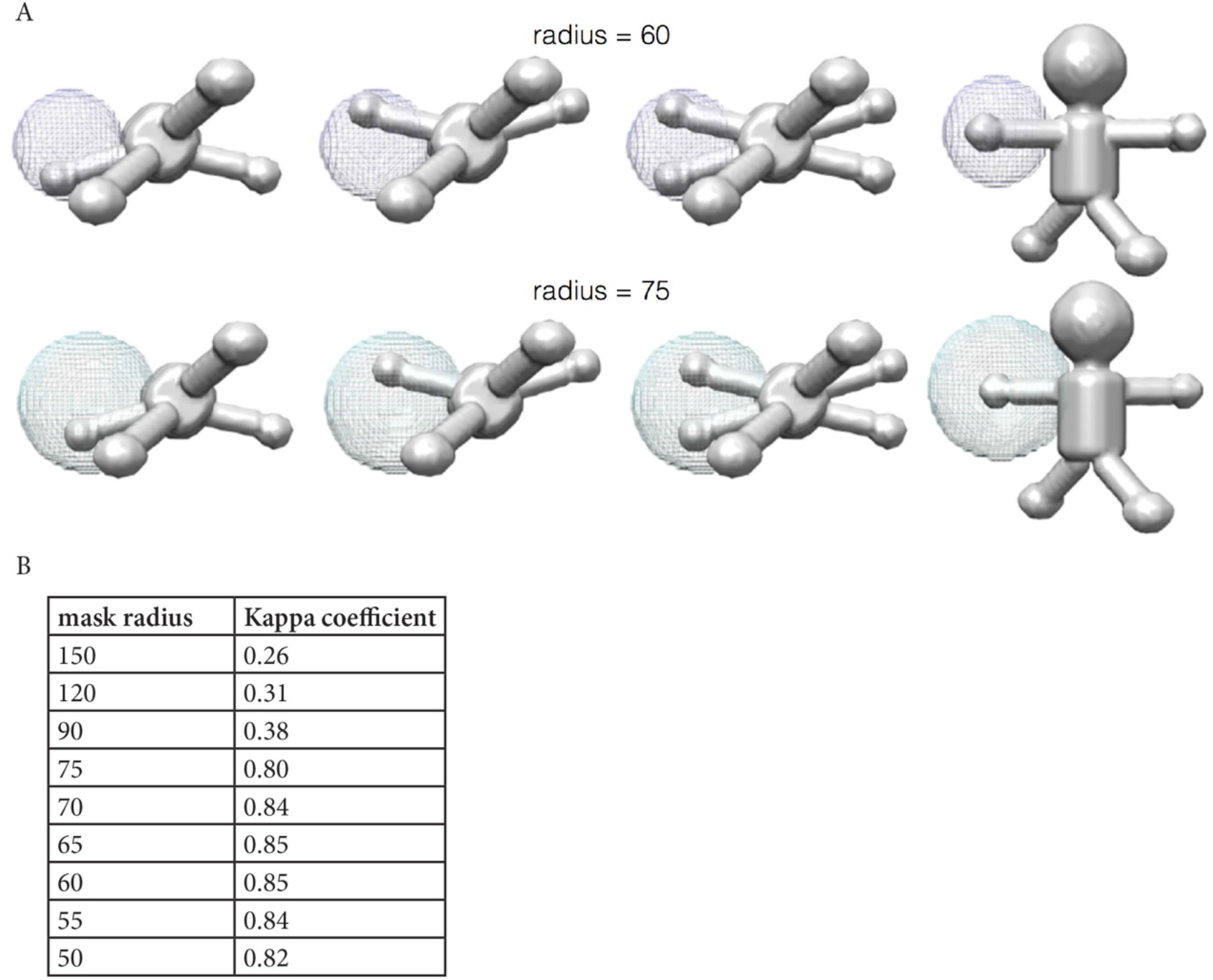
titration of mask size used for focused classification. Focused classification parameters could be tuned for optimal performance with this particular dataset. Here, the mask size was varied, and the results were followed by monitoring κ. (A) Two different mask sizes are displayed, applied to an asymmetric unit around the arms/hands. (B) The results of focused classification with different mask radii. Here, a 60 Å mask performs optimally, which effectively represents a tight mask that completely encompasses only the mobile area.

**Supplementary Figure 4 -.**
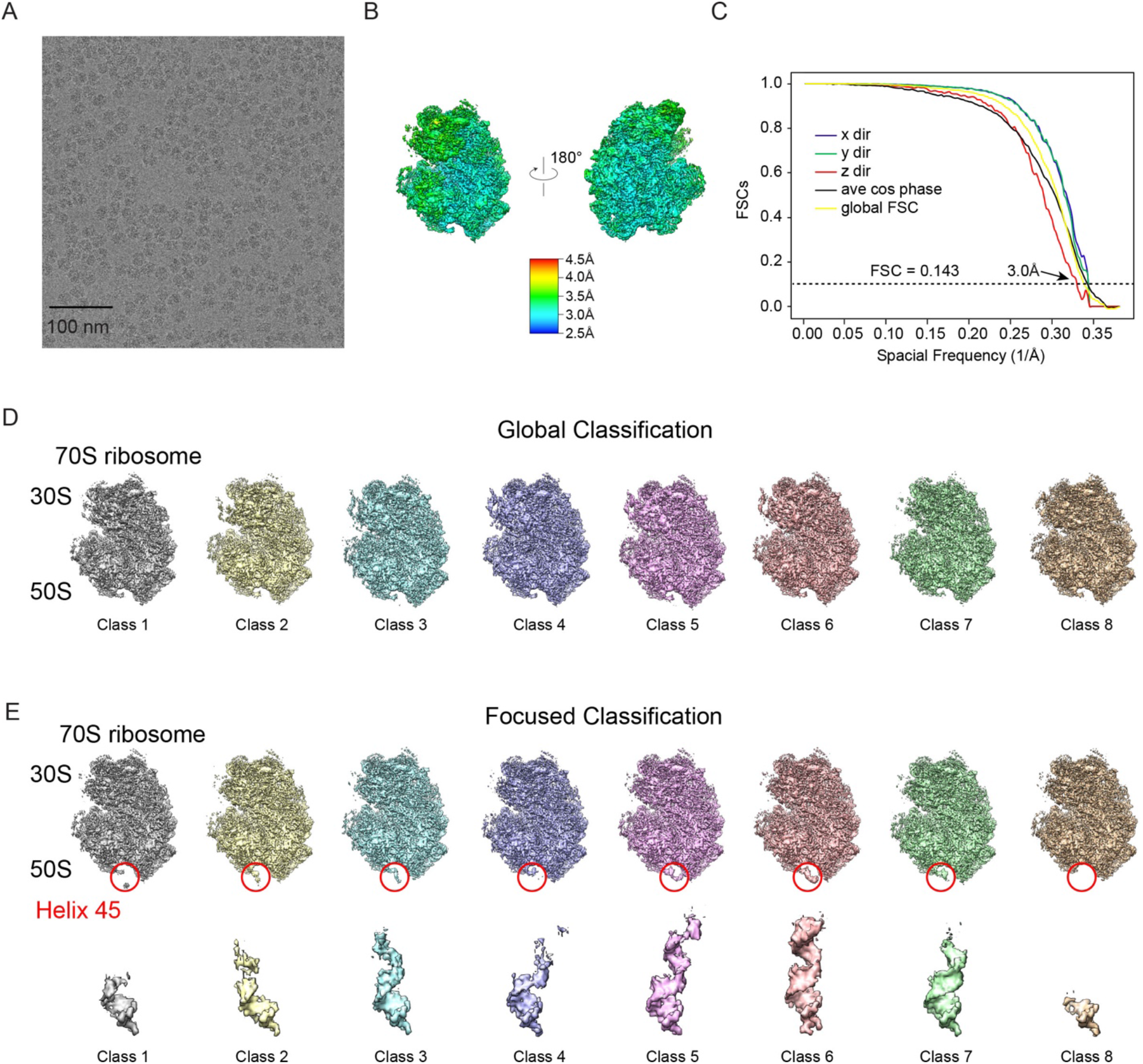
Cryo-EM data for HIV-1 TAR—ribosome fusions. (A) Example raw image collected for TAR-labeled ribosomes. (B) Initial single-model refinement, colored by local resolution and (C) the corresponding FSC curves. (D) Classes generated from global 3-D classification showing a lack of density in the region of helix 45. (E) Classes from focused 3-D classification, with the mask applied to the region of TAR fusion, denoted by a red circle with the corresponding densities of the TAR hairpin in the absence of the ribosome scaffold below.

